# Plasticity, geographic variation and trait coordination in blue oak drought physiology

**DOI:** 10.1101/2023.08.20.553748

**Authors:** Leander D.L. Anderegg, Robert P. Skelton, Jessica Diaz, Prahlad Papper, David D. Ackerly, Todd E. Dawson

## Abstract

Our ability to predict drought stress across the landscape remains limited. This uncertainty stems in part from an incomplete understanding of within-species variation in hydraulic physiology, particularly coordinated variation across multiple traits. This variation reflects genetic differentiation among populations (ecotypic variation) and phenotypic plasticity. We examined among-population differentiation in morphological and hydraulic traits in California blue oak (*Quercus douglasii*) using a 30 year-old common garden. We compared trait differentiation and trait-trait coordination in the garden to wild phenotypes from the original source populations. We found remarkably limited among-population differentiation in all traits in the common garden but considerable site-to-site variation in the field that could rarely be explained with site climate variables. Trait-trait relationships were also stronger in the field than in the garden, particularly links between leaf morphology, leaf hydraulic efficiency and stem hydraulic efficiency. Only four trait-trait relationship were present in both the wild and garden, but 12 of 45 relationships showed significant wild phenotypic correlations, with strong coordination among leaf and stem hydraulic efficiency apparently mediated by leaf size. Ultimately, we found limited evidence for ecotypic variation but considerable geographic in phenotypic integration in the wild, suggesting considerable acclimation potential in the face of climate change.

## Introduction

In the 21^st^ century, trees living in a hotter, more variable and often drier world will need to acclimate or adapt to avoid local extirpation. Recent drought- or heat-induced forest mortality events highlight the vulnerability of even highly drought-adapted forests to climate change (Allen *et al*. 2010; Brodribb, Powers, Cochard & Choat 2020; Hammond *et al*. 2022). Unfortunately, despite multiple decades of concerted effort to understand the causes of drought-induced forest mortality, we still struggle to predict when and where trees will die off during drought (Trugman, Anderegg, Anderegg, Das & Stephenson 2021).

Part of the large uncertainty about drought vulnerability is a poor understanding of physiological variation within species (Trugman *et al*. 2021). Within-species variation can be substantial (Martinez-Vilalta *et al*. 2009; Mclean *et al*. 2014; Anderegg *et al*. 2021) and can decrease climate vulnerability in some populations (Laforest-Lapointe, Martinez-Vilalta & Retana 2014; Garcia-Forner, Sala, Biel, Savé & Martinez-Vilalta 2016). Drought resistance is the complex result of numerous plant traits, all of which potentially vary across populations within a species. Xylem resistance to embolism, often characterized by P50 or the water potential inducing 50% xylem embolism, has been identified as a key drought tolerance trait (i.e. a trait that allows a plant to experience drought stress without suffering physiological damage, (Levitt 1980; Anderegg & HilleRisLambers 2016)), that mediates mortality differences among species (Anderegg *et al*. 2016). However, within individual species, P50 is often quite consistent across populations (Lamy *et al*. 2013). For instance, in blue oak (*Quercus douglasii* Hook & Arn), a culturally and ecologically important tree species that dominates many of the oak savannas in California, USA, Skelton *et al*. (2019) previously found no evidence for local adaptation or plastic differences in leaf or stem P50 across blue oak’s geographic range. This implies that the threshold for hydraulic damage is remarkably consistent across blue oak populations, despite large geographic differences in water availability (Fig 1).

**Figure 1:**
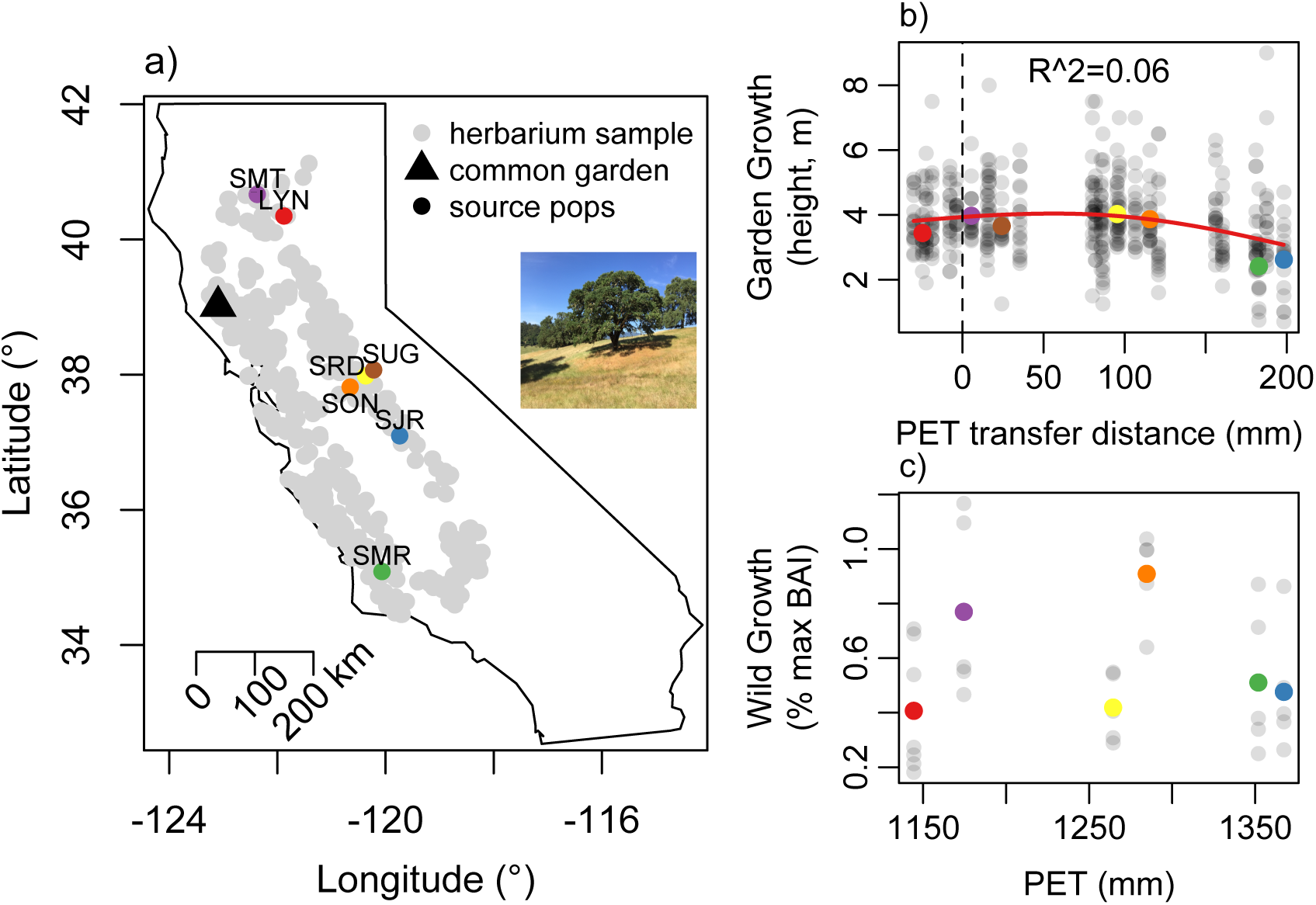
(a) Geographic distribution of blue oak (Quercus douglasii Hook & Arn) based on herbarium specimens, with focal source populations and common garden location. (b) Height growth in the common garden (gray points show all individuals from full 26 populations, colored points show population means of seven focal populations) as a function of the potential evapotranspiration (PET) ‘transfer distance’ (difference between garden and source locations). Garden growth shows little evidence of home-site advantage, with only subtle among-population growth differences and maximum growth reached at positive transfer distances. (c) Wild growth as % of maximum Basal Area Increment (BAI) standardized for tree size as a function shows no significant relationship with site climate variables (plotted against PET for illustration).

How, then, do species such as blue oak adjust physiologically to deal with geographic variation in drought stress if their tolerance is fixed? The answer likely lies in coordinated adjustments of numerous traits, particularly drought avoidance traits (traits that allow a plant to avoid experiencing drought stress under water deficit). For example decreasing leaf area to sapwood area ratio (A_L_:A_S_) or increasing tissue-specific hydraulic efficiency can help plants avoid drought stress by minimizing the water potential drop required to maintain transpiration fluxes (Gleason, Butler, Waryszak & Graham 2013; Mencuccini *et al*. 2019). Previous studies have found conflicting patterns of trait coordination within species, ranging from consistent increases in tissue robustness (leaf mass per area and wood density) and decreases A_L_:A_S_ in dry populations of multiple Australian tree species (Anderegg *et al*. 2021) to extremely unpredictable patterns of trait coordination within various Iberian tree species (Rosas *et al*. 2019). Indeed, the extent to which physiological and morphological traits are coordinated within species across different environments remains poorly understood (Blackman, Aspinwall, Resco de Dios, Smith & Tissue 2016). It remains an open question how individual species coordinate variation across many traits to develop drought resistant phenotypes in dry places, as broad evolutionary patterns of trait covariation among species often break down within species (Anderegg *et al*. 2018; Messier, Violle, Enquist, Lechowicz & McGill 2018; Rosas *et al*. 2019).

Equally important to the total amount of within-species variation are the drivers of this variation. Intraspecific variation can arise from two distinct processes with drastically different implications for near-term climate responses. First, genetic differentiation among populations can drive spatial phenotypic variation, caused be either neutral processes (e.g. drift, founder effects) or spatio-temporal variation in natural selection on heritable fitness-related phenotypes that leads to local adaptation of functional traits (Alberto *et al*. 2013). Substantial geographic variation in drought physiology due to genetic differentiation among populations would imply that certain populations or genotypes (and not others) are drought resistant and that gene-flow (assisted or natural) is necessary for other populations to manifest improved drought resistance (Hoffmann, Miller & Weeks 2021). Population-level genetic differentiation is widespread in plants, particularly species with large population sizes (Leimu & Fischer 2008; Savolainen, Lascoux & Merilä 2013). Indeed, provenance trials or common gardens (where multiple populations of a species are planted in a common environment) often detect genetic differentiation in performance (e.g. growth) among populations in trees (Alberto *et al*. 2013; Ramírez-Valiente, Santos del Blanco, Alía, Robledo-Arnuncio & Climent 2022). While phenotypic differences among populations growing in a common environment cannot demonstrate local adaptation *per se* (i.e. that among-population genetic variation is adaptive), traits showing among-population genetic differentiation that is related to the climate of the source population can highlight phenotypes that may show the fingerprint of past selection (Schwinning, Lortie, Esque & DeFalco 2022).

Alternatively, trait plasticity, or the ability of a genotype to manifest a broad range of phenotypes depending on environmental cues, can generate geographic variation in water stress-related traits (Blackman, Aspinwall, Tissue & Rymer 2017). If plasticity drives geographic trait variation (and plasticity itself does not differ among populations), all populations of a species could potentially manifest drought resistant traits given the right environmental cues. As sessile organisms that experience a range of environments over their lifetime and even in different parts of the same organism (e.g., light availability in different parts of the canopy), plants show marked plasticity in many traits (Poorter, Niinemets, Poorter, Wright & Villar 2009; Palacio-López, Beckage, Scheiner & Molofsky 2015; Keenan & Niinemets 2016). Indeed, substantial plastic responses to drought and cold stress have been observed in multiple tree species (Gimeno, Pías, Lemos-Filho & Valladares 2009; Gárate-Escamilla, Hampe, Vizcaíno-Palomar, Robson & Benito Garzón 2019).

We quantified the extent and drivers of geographic variation in drought-related traits within mature blue oak trees. Recent seedling experiments in blue oak (B. McLaughlin et al., 2022; B. McLaughlin et al. in prep) and other oak species (Ramírez-Valiente & Cavender-Bares 2017; San-Eufrasio *et al*. 2020) have revealed genetic variation in the performance of seedlings under drought related to their climate of origin. However, the extend of ecotypic variation in adult oaks remains largely unknown due to the difficulty of maintaining common garden experiments. Meanwhile, blue oak has also shown substantial plastic changes in leaf morphology during severe drought (Weitz 2018), hinting at its large potential for phenotypic plasticity. Thus, we investigated how blue oak, which grows across a huge range of water availability (Fig 1), manifests phenotypic variation across the landscape. This could manifest as geographic variation in drought avoidance traits such as hydraulic efficiency and leaf area:sapwood area ratios, as well as variation in drought tolerance traits beyond P50, such as leaf robustness (specific leaf area or SLA, leaf dry matter content or LDMC). We then attempt to disentangle the extent of among-population genetic differentiation versus plasticity in this phenotypic variation and trait coordination.

Specifically, we asked 1) how much do drought-related traits vary across the landscape, 2) in a 30 year old, reproductively mature common garden, is there evidence for among-population genetic variation underlying this landscape trait variation and 3) how is drought resistance coordinated across different traits and tissues, in common garden and wild populations? We sought to disentangle the relative roles of population-level differentiation versus plasticity by measuring drought-related traits in a common garden experiment compared to the traits of the source populations in the wild. In the garden, trait differences between populations are indicative of between-population genetic differentiation (genotype or G effects, assuming limited non-genetic carryover such as maternal effects). Meanwhile, variation measured in the wild source populations represents a combination of genetic and plastic effects (G + environment or E effects + genotype-by-environment or GxE effects). Thus, the difference between trait variation in the common garden (G only) and the wild (G + E + GxE) can provide some inference about the extent of plasticity.

## Methods

Blue oak (*Quercus douglasii* Hook & Arn.) is a deciduous species, which is endemic to California but widespread and abundant throughout the state in low elevation woodlands around the California Central Valley (Fig. 1, Stahle et al., 2013). Blue oak experienced substantial and spatially widespread mortality during a major drought from 2012-2016, particularly in the southern portion of its geographic range (Brown, McLaughlin, Blakey & Morueta-Holme 2018; McLaughlin *et al*. 2020). This mortality highlights the potential vulnerability of blue oak in a changing climate, necessitating an improved understanding of blue oak drought tolerance in space and time to support proactive oak management. We built on the work of Skelton et al. (2019) on xylem vulnerability to embolism to quantify how morphological and hydraulic traits of leaves and stems beyond P50 vary across seven populations of blue oak in the wild and in a ∼30 year-old, reproductively mature common garden. These populations were selected to span the geographic range and range of moisture availability of *Q. douglasii* (Fig. 1, Table S1), using only populations whose original acorn source (for the common garden) had been accurately relocated, and whose source populations were accessible based on current land tenure.

### Climate data

Climate data for the source populations, both 1951-1980 climate normals and 2017-2018 water year meteorology (the year of trait sampling), were extracted from the California state-wide Basin Characterization Model (Flint, Flint, Thorne & Boynton 2013). Only historical source population climate normals were used for garden analysis, while historical climate, sampling year meteorology, and the sampling year anomaly (sampling year – 30yr normal) were used for analysis of wild traits.

### Common garden sampling

To investigate the amount of among-population genetic trait differentiation, we sampled an existing provenance trial/common garden planted at the Hopland Research and Extension Center (CA, USA), a fairly mesic site for blue oak (Figure 1). This trial was planted in 1992 with acorns collected from 26 *Q. douglasii* populations across California and planted in a randomized block design (J. McBride, pers. comm.; see also McBride *et al*. 1997). One year-old seedlings were planted into the garden in six replicated blocks with nine sub-replicate individuals per population per block. Individuals were grown from bulked acorns collected from 10 maternal trees per population, and thus represent some combination of half sibs and more distant relatives (families were not tracked at planting). In 2001-2002, six of the nine sub-replicate individuals per population were thinned in each block to reduce competition. By 2018, mean tree height was 3.79 m, and 13 of the 468 planted individuals had died (Papper & Ackerly 2021). Due to the time-intensive nature of hydraulic phenotyping, we subsampled seven populations, specifically selected to capture as much of *Q. douglasii’*s aridity range as possible (Fig. 1). For each of the seven populations, we sampled five to six individuals (one per block in 5-6 blocks) for hydraulic traits, leaf traits and branch wood density. Between late-April and mid-June of 2018, we collected one to three >1 m long branches to subsample for hydraulic measurements from the sunlit, south-facing portion of the canopy in the early morning (before 9 am local time). We relaxed the xylem by repeatedly recutting the stem underwater, and then placed the branches in large plastic bags for immediate transport back to the lab. One to two individuals from each of the seven sampled populations were collected from the garden at each sampling point (once or twice weekly) to control for temporal variation in hydraulic traits. Three small terminal twigs and subtending leaves were collected at the same time for the measurement of morphological traits. Tree diameter at 50 cm and total tree height was also surveyed for every common garden tree in the winter of 2017 and used to calculate the total stem volume of the largest stem (assuming the stem was a cylinder) as a metric of total growth rate since planting.

### Wild population sampling

We collected branches for trait measurement from six to eight mature trees from the source populations used in the garden. The exact maternal trees from which acorns were collected for the common garden could not be located, so the selected wild trees represented a sampling of reproductively mature, visibly healthy trees near the documented acorn collection location. Within a location, individuals were 5-50 m apart, often including all large trees present within a 500 m search area around the original collection site. Minimal ongoing or historic tree mortality (e.g., dead snags or stumps) were noted at the source populations and all wild trees were at least 30 yrs old, making adaptive evolution since seed collection unlikely. Large branches (typically >1.5 m long and >4 cm basal diameter) from the south-facing canopy were collected in the same manner as in the common garden. Short tree cores were also collected from the sampled trees for calculation of radial growth rates for the past decade. Sites were sampled between late April and early June 2018, roughly controlling for time since leaf out (i.e. earlier leafing sites were collected first so that all sites could be collected after full leaf expansion, ∼1-2 months after leaf flush).

### Morphological Traits

We selected three terminal branches from the south-facing sun-exposed canopy for morphological measurements (Table 1), cut at the prior year bud scar and including all current year stem and leaf tissue. Branches were rehydrated for >12 hrs using the ‘partial rehydration’ method (Pérez-Harguindeguy *et al*. 2013). We then measured specific leaf area (SLA, cm^2^ wet leaf area per g^−1^ leaf dry mass), leaf dry matter content (LDMC, g leaf dry mass per g^−1^ leaf wet mass), mean leaf size (cm^2^ fresh leaf area), leaf dry mass to stem dry mass ratio (M_L_:M_S_), and leaf area to sapwood area ratio (A_L_:A_S_, cm^2^ leaf area per mm^2^ stem area underneath bark, or the inverse of the Huber Value) on these current year terminal twigs and all of their subtending leaves using calipers, balances, and ImageJ image analysis software. We calculated mean leaf size as the average area per leaf for each terminal branch. We also measured wood density (WD, g dry mass per cm^3^ wet volume) on one to five branch disks 2-5 cm in diameter cut from the basal end of branches collected for hydraulic sampling. Bark was removed from branch disks and wet volume was measured via water displacement on a balance.

**Table 1:** Trait definitions and abbreviations as employed by this study.

|  |  |
| --- | --- |
| Trait | Definition |
| SLA | Specific leaf area ($\text{cm}^2 \text{g}^{-1}$) |
| LDMC | Leaf dry matter content (g dry mass per g fresh mass) |
| WD | Wood density |
| M<sub>L</sub>:M<sub>S</sub> | Terminal branch leaf dry mass to stem dry mass ratio (g/g) |
| A<sub>L</sub>:A<sub>S</sub> | Terminal branch leaf area to sapwood area ratio ($\text{cm}^2 \text{mm}^{-2}$) |
| Leaf size | Average area of a single leaf based on all leaves on a terminal branch ($\text{cm}^2$) |
| k<sub>leaf</sub> | Leaf conductance per unit leaf area ($\text{mmol m}^{-2} \text{s}^{-1} \text{MPa}^{-1}$) |
| k<sub>stem</sub> | Leaf area-specific stem conductance of full current year terminal branch ($\text{mmol cm}^{-2} \text{s}^{-1} \text{kPa}^{-1}$) |
| K<sub>s</sub> | Sapwood area-specific stem conductivity per branch length ($k[\text{stem}] * A_L:A_S * \text{length}$, $\text{mmol cm}^{-1} \text{s}^{-1} \text{kPa}^{-1}$) |
| P50<sub>leaf, stem</sub> | Leaf and stem P50 measured with optical technique from Skelton et al. 2019 |
| Growth | **Garden:** stem volume attained at 30 yrs, based on basal diameter and height of largest stem assuming cylindrical stem.<br>**Wild:** percent of size-specific maximum basal area increment (based on rangewide 90<sup>th</sup> percentile BAI at a given DBH) |

For all analyses using individual or population average traits, we also included leaf and stem P50 values (the xylem tension causing 50% embolism) from Skelton *et al*. (2019), which were measured on branches collected at the same time and using the same methods as branches collected for hydraulic efficiency measurements reported here. However, P50 measurements had few or no replicates per individual and were thus not suitable for branch-level analyses.

### Hydraulic traits

Due to extremely long vessel lengths in oaks (often >1 m in branches of *Q. douglasii*, based on pressurized branches recut at the basal end until air bubbles were seen at the distal end; RPS and LDLA *pers. obs.*), classic hydraulic methods using stem segments were impossible without considerable open vessel artifacts. Instead, we performed hydraulic measurements on terminal branches of 1-2 mm basal diameter using the vacuum chamber method (Kolb, Sperry & Lamont 1996). We targeted current year growth when possible (similar to branch sampling for morphology), but included up to three years of growth when the current year was <2 cm long and stem diameter was not sufficiently large to fit in hydraulic tubing. For each terminal branch, all subtending leaves were cut under water at the petiole with a razor blade, and the entire stem was inserted into a vacuum chamber. Flow from a scale, through the branch and out the cut petioles was induced by subjecting the stem to a ∼60 kPa vacuum. Nevertheless, any stem that was suspected of having an open vessel (i.e. any stem that had a high apparent conductance) was checked for open vessels by attaching to pressurized nitrogen for ∼10 minutes and checking under water for air bubbles from the stem or petioles. The leaf area of all subtending leaves was measured via flatbed scanner and ImageJ (Schneider, Rasband & Eliceiri 2012), basal stem diameter calculated from 4 radii, and the length of the stem and all branches measured with digital calipers.

Raw stem conductance was standardized two different ways. First, raw conductance was standardized per unit leaf area, here termed k_stem_, indicating leaf area-specific stem conductance (i.e. not standardized by path length, so including the effects of differential stem growth in long versus short stems, Table 1). Raw conductance was also divided by stem cross-sectional area and multiplied by total stem length (including length of branches if stem was branched) to produce sapwood area specific conductivity, or K_s_.

Leaf hydraulic conductance was also measured on terminal branches (cut from the same >1 m long branch as stem conductance) using the ‘rehydration kinetics’ (RK) method (Brodribb & Holbrook 2003). Because leaves often had very small and irregular petioles, we measured multiple leaves attached to a terminal twig. Leaf conductance was an order of magnitude lower than stem conductance, meaning the effect of the stem in the conductance observed via the RK measurements was negligible. Total leaf area for each sample was calculated via flatbed scanner and ImageJ for each sample, and leaf-area specific leaf conductance, k_leaf_ was calculated. Only measurements from leaves with an initial water potential < −0.2 MPa and > −2 MPa were analyzed to avoid large measurement errors in samples with small pressure gradients or potential embolism. Skelton et al. (2019) measured leaf and stem xylem vulnerability to embolism on the same individual using the optical method. Large (>1m) branches for vulnerability curves were also collected in the spring of 2018, often on the same day as for hydraulic measurements reported here. We include the P50 measurements (xylem potential causing 50% embolism) from Skelton et al. 2019 for comparison with the traits measured here, but note that replication was generally much smaller for P50 (1-2 replicates per individual as opposed to 3+ for all traits reported here) so analyses could only be performed at the individual rather than replicate level for P50.

### Growth rate

We calculated growth rates for the common garden individuals based on a survey of tree diameter at 50 cm and stem height during the winter of 2017, when trees were approximately 27 years old. We tested for evidence of local adaptation by modeling height growth as a function of ‘transfer distance’ between the source location and the common garden using generalized additive mixed models penalized spline smoothers for climate predictors (gamm function from the *mgcv* package in R) with population as a random intercept. A concave down relationship between growth and transfer distance that peaks at transfer distances near zero would indicate ‘home site advantage’ or that populations from climates most similar to the garden climate perform best in the garden, though the fact that the garden was at the mesic edge of blue oak’s distribution decreases our power to detect this peak in transfer distance. 30 yr climate normals for the seed sources were used as potential predictors and the best fit model was selected using AIC. Results were qualitatively similar with height and 50 cm diameter, so only height is reported.

Growth rates were calculated for the wild trees from tree cores, which were collected via increment borer at 1.3 m height; cores were mounted and sanded, and 5 years worth of rings identified using a dissecting microscope. The length of the 2013-2018 5-year growth period was measured via digital calipers and radial growth was converted to Basal Area Increment (BAI) based on tree DBH. Tree BAI is a function of tree size (Fig. S1), so BAI was standardized for tree size by calculating the percent of size-specific maximum Basal Area Increment (BAI) over the 2013-2018 period. Size-specific maximum BAI (the fastest rate a tree of a given DBH was observed to grow) was calculated for each tree’s DBH based on the 90^th^ quantile regression of BAI versus tree DBH for all study trees (n=32) plus tree core data from 29 trees spanning five additional sites across California to expand the range of sampled tree sizes (total DBH range 7.8cm-104cm, Fig. S1). The observed BAI was then divided by the maximum BAI for each tree to produce ‘% max BAI’. Tree cores were collected in Oct of 2018, but could not be collected from one site due to loss of site access (n=6 sites rather than 7).

### Variance decomposition analysis

We performed variances decompositions, separately in the garden and the wild, on all branch-level trait measurements to quantify trait variation within canopy (among branches), among individuals within populations, and among populations. We fit linear mixed-effects models separately for the garden or the wild traits, with a fixed intercept term, a random intercept for population and a random intercept for individual nested within population. We employed the lmer() function from the *lme4* (v 1.1-28) and *lmerTest* (v 3.1-3) packages. We then extracted the random effect variance parameters, and calculated the proportion of total trait variance that was attributed to among-population differences (population random effect), individual differences within populations (individual random effect) or variation within individual tree canopies (residual variance). We also calculated the coefficient of variation (CV = trait mean / trait standard deviation) for each trait in the garden and in the wild. Finally, to compare the total trait variation in the garden versus in the wild while accounting for unequal sample sizes, we bootstrapped variance estimates, randomly sampling with replacement each dataset 1000 times with sample number set to the minimum sample size of the two datasets (wild versus garden) for each trait. We then compared the median bootstrapped trait variance for the wild versus garden.

### Among-population trait differentiation and climatic variation

We tested for significant among-population trait differences by averaging trait values per individual and then performing one-way ANOVAs separately in the garden and in the wild, and calculated the omega-squared (ρο^2^) as an estimate of the proportion of variance explained by among-population differences. Garden block was never a statistically significant factor in the garden, and was excluded from the final ANOVAs (fitted using the aov() function in the *stats* package, v 4.1.2). We then used AICc (Akiake’s Information Criterion corrected for small sample sizes, AICc() function from the *MuMIn* package, v 1.43.17) to select the best single climatic predictors of among-population differences for each trait. First, we determined whether a random intercept for population was required to account for non-independence among individuals in a population by fitting an ‘over the top’ fixed effects structure (Zuur, Ieno, Walker, Saveliev & Smith 2009), including mean annual precipitation, mean annual potential evapotranspiration and mean minimum temperature, and using likelihood ratio test to determine whether the population random effect was needed (using the gls() and lme() functions from the *nlme* package, version 3.1-155). We then used AICc to select the best single climate predictor for each trait using linear models (lm in the *stats* package) or linear mixed models with a population random intercept (lmer in the *lme4* package) and assessed significance of the best predictor using t-tests, using Satterthwaite’s degrees of freedom from the *lmerTest* package (Kuznetsova, Brockhoff & Christensen 2016). For the common garden, potential individual predictors included only source population 30 year climate normals (1951-1980) of mean annual precipitation (PPT_30yr_), mean annual potential evapotranspiration (PET_30ry_), mean annual actual evapotranspiration (AET_30yr_), mean annual climatic water deficit (CWD_30yr_=PET_30yr_-AET_30yr_), and mean annual minimum temperature (Tmin_30yr_). For wild traits, potential univariate predictors included historic 30 year climate normal as well as growth year (2017-2018) weather information: wet season (Nov-May) minimum temperature (Tmin_gy_), maximum temperature (Tmax_gy_), precipitation (PPT_gy_), potential evapotranspiration (PET_gy_), actual evapotranspiration (AET_gy_) and climatic water deficit (CWD_gy_), as well as the anomaly of the 2018 water year from the site 30 year normal (PPT_anom_, PET_anom_, AET_anom_, CWD_anom_). We visually examined quantile-quantile plots and other patterns in the residuals to identify outliers and ensure model assumptions were met.

### Trait-trait coordination

To examine the magnitude of phenotypic (genetic plus plastic variation in the wild) and genetic (variation in the garden) trait correlations, we calculated Pearson’s correlation coefficients for all pairs of individual-averaged traits in the wild and in the common garden. We visualized the correlation structure using the corrplot() function in the *corrplot* package (v 0.92). We then used a similar procedure to the bootstrapped variance comparisons to test for significant differences between genetic and phenotypic trait correlations, randomly sampling individuals with replacement from the entire wild and entire common garden datasets to produce two populations of equal size to the smallest of the garden or wild complete pairwise observations, resampling 1000 times per trait combination. We then calculated the difference between the trait-trait correlations in each bootstrapped pair of garden and wild datasets and determined the two-tailed probability that the distribution of differences did not include 0 (e.g., alpha < 0.05 if either the 2.5^th^ percentile of differences was greater than zero or the 97.5^th^ percentile of differences is less than zero, alpha < 0.1 if the 5^th^ and 95^th^ percentile range did not include zero, and alpha < 0.2 if the 10^th^ to 90^th^ percentile range did not include zero).

All code for analyses and figure generation can be found at https://github.com/leanderegg/BlueOakGarden.git and all data are available on Dryad at [insert Dryad link upon acceptance](Data citation). *NOTE all data are currently in the github repository for peer review

## Results

Growth in the common garden showed relatively small among-population variation and little evidence of local adaptation. The historic PET of the seed source best explained among population differences in garden growth (using heights of all surviving individuals including all 26 original source populations), but only explained a small fraction of the variation in growth (R^2^=0.05 for linear model, Table S2; R^2^ = 0.06 for penalized regression spline, Fig. 1b). The relationship between climate transfer distance and growth was concave down but growth peaked at positive transfer distances (higher source PET than the garden), demonstrating little ‘home site advantage’ that would indicate local adaptation (Fig. 1b). Meanwhile, growth in the wild differed drastically among populations but was not strongly related to any climate variable (Table S2, Figure 1c). Source environmental conditions did not appear to drive garden growth through parental stress, as growth in the wild was uncorrelated with growth in the garden (Fig. S2).

Genetic variation among populations was low in all morphological, allocation and hydraulic traits in the common garden (Fig. 2a). However, all traits except for stem sapwood area-specific hydraulic conductivity (K_s_) and leaf and stem vulnerability to embolism (P50_leaf_, P50_stem_) showed substantial among-population variation in the wild. Among-population variation in the wild was consistently 25%-30% of total trait variation in most traits, with SLA and LDMC showing even larger among-population variation (46% and 40%, respectively, Fig. 2b). Meanwhile, branch-to-branch variation within tree canopies consistently made up the majority of trait variation in the common garden, constituting >50% of total variation in all traits except WD (42% of total) and over 75% of total variation in k_leaf_, k_stem_, and K_s_ (Fig. 2a). In the wild, within-tree variation was typically less than 50% of total variation, except in M_L_:M_S_ and the three hydraulic traits (Fig. 2b).

**Figure 2:**
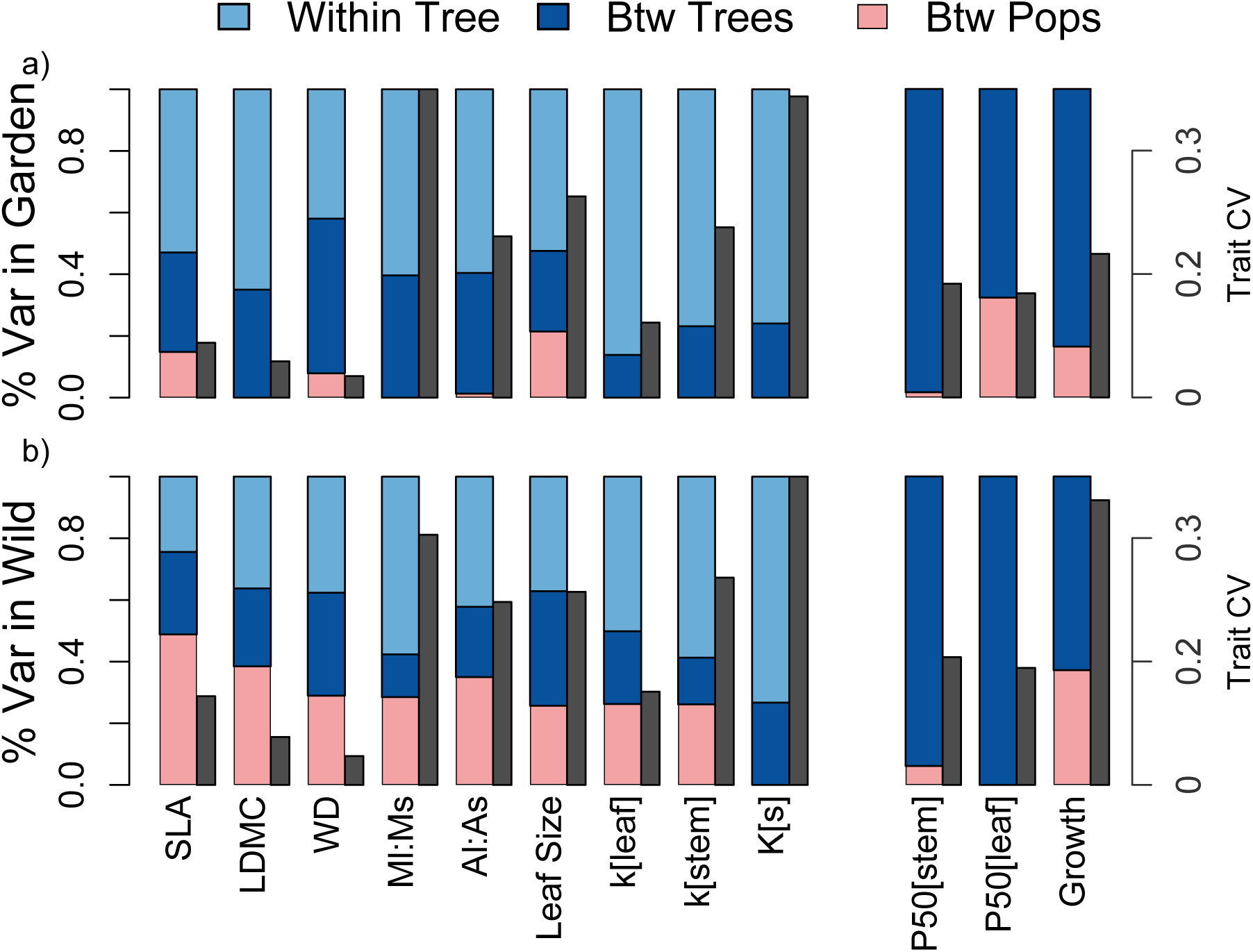
Variance decomposition of all branch-level measurements in the common garden (a) and wild source populations (b). Grey bars beside the colored bars show the magnitude of the trait coefficient of variation. P50_stem_, P50_leaf_ and Growth do not have replicates within tree, so only contain between tree and between population variance components.

When traits were averaged to the individual, 40-60% of the total tree-to-tree variation in almost all traits was between populations in the wild (Fig. 3a) and population differences were significant in 8 of 11 traits (all but K_s_, P50_leaf_, and P50_stem_) plus growth (ANOVAs p <0.05, Table S2). Meanwhile, typically <30% of tree-to-tree variation was between populations in the garden, and population differences were only significant in 2 of 10 traits (SLA and leaf size) plus growth in the garden (Table S2). Leaf size was the one notable trait that showed significant and similar among-population differentiation in both the wild and garden.

**Figure 3:**
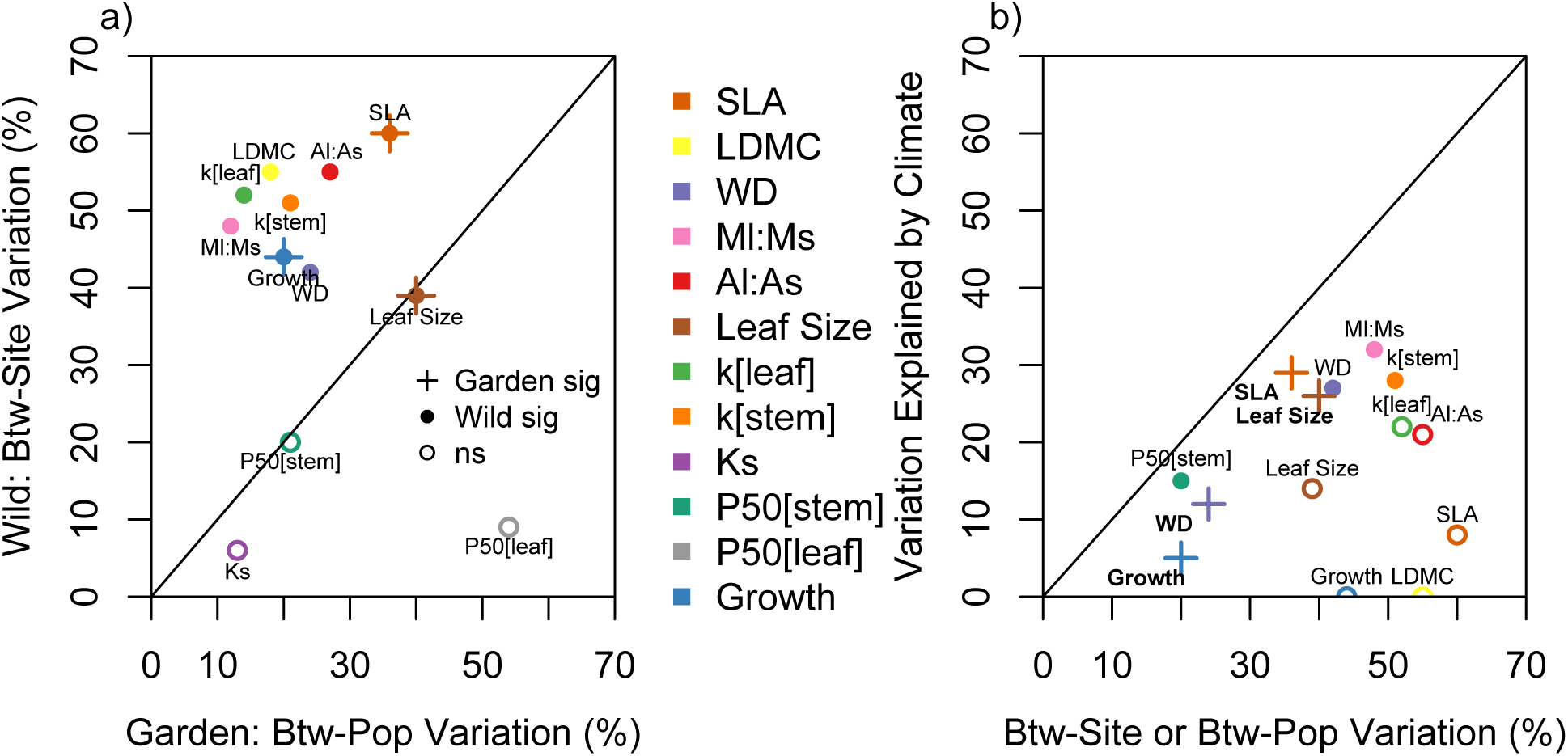
(a) % of the total tree-to-tree variation attributable to population in the garden versus attributable to sites in the wild. Traits with significant between-site variation (ANOVA p <0.05) in the wild are shown as filled points, while traits with significant between population variation in the garden are shown with crosses. (b) The among-population and among-site variation that could be explained with climate variables, including only traits with significant among-population variation (ANOVA p<0.05). Y-axis shows R^2^ of best climate predictor based on AICc, filled points indicate predictors that are statistically significant for wild traits, open points show traits with significant among-population differentiation but non-significant climate predictors, and crosses with bold text labels show garden traits with significant climate predictors (Table S2). Black line shows 1:1 relationship.

However, despite the consistent among-population variation in wild traits, climate predictors (historical climate normals, meteorology of the water year of sampling, or the sampling year anomaly from historical climate) very rarely explained differences among populations (Fig. 3b). Based on AICc of linear or linear mixed models (when there was support for a site random intercept, see Table S2), univariate climate predictors were only statistically significant for four wild traits (Fig 3b, Fig S3,S4, Table S2) with M_L_:M_S_ decreasing and k_stem_ increasing with Tmin_gy_, P50_stem_ decreasing (growing more negative) with AET_30yr_, and WD decreasing with AET_anom_ (though the relevance of annual anomaly for wood density of 5+ year old branches is suspect). Meanwhile, in the garden three traits and growth were significantly associated with source population climate (Table S2). In the garden, increasing PPT_30yr_ was significantly associated with decreasing wood density and increasing leaf size, while increasing PET_30yr_ was significantly associated with increasing SLA (Fig S3,S4).

The among-population variance was smaller in the garden than the wild for all traits except leaf size (Fig. 4). However, the total among-individual variance was usually smaller in the wild than the garden (garden:wild ratio <1), but only significantly so (alpha <0.05) for SLA, LDMC, and WD based on bootstrapped variance estimates. Meanwhile, M_L_:M_S_, k_leaf_ and k_stem_ had similar total variances in the wild and the garden despite much larger among-population variance in the wild. K_s_, P_50[stem]_ and P_50[leaf]_ had similar variances in the wild compared to the garden and no significant among-population differentiation in either location (Table S2), while growth could not be compared because of differing metrics (% of max basal area increment vs 30 yr stem volume).

**Figure 4:**
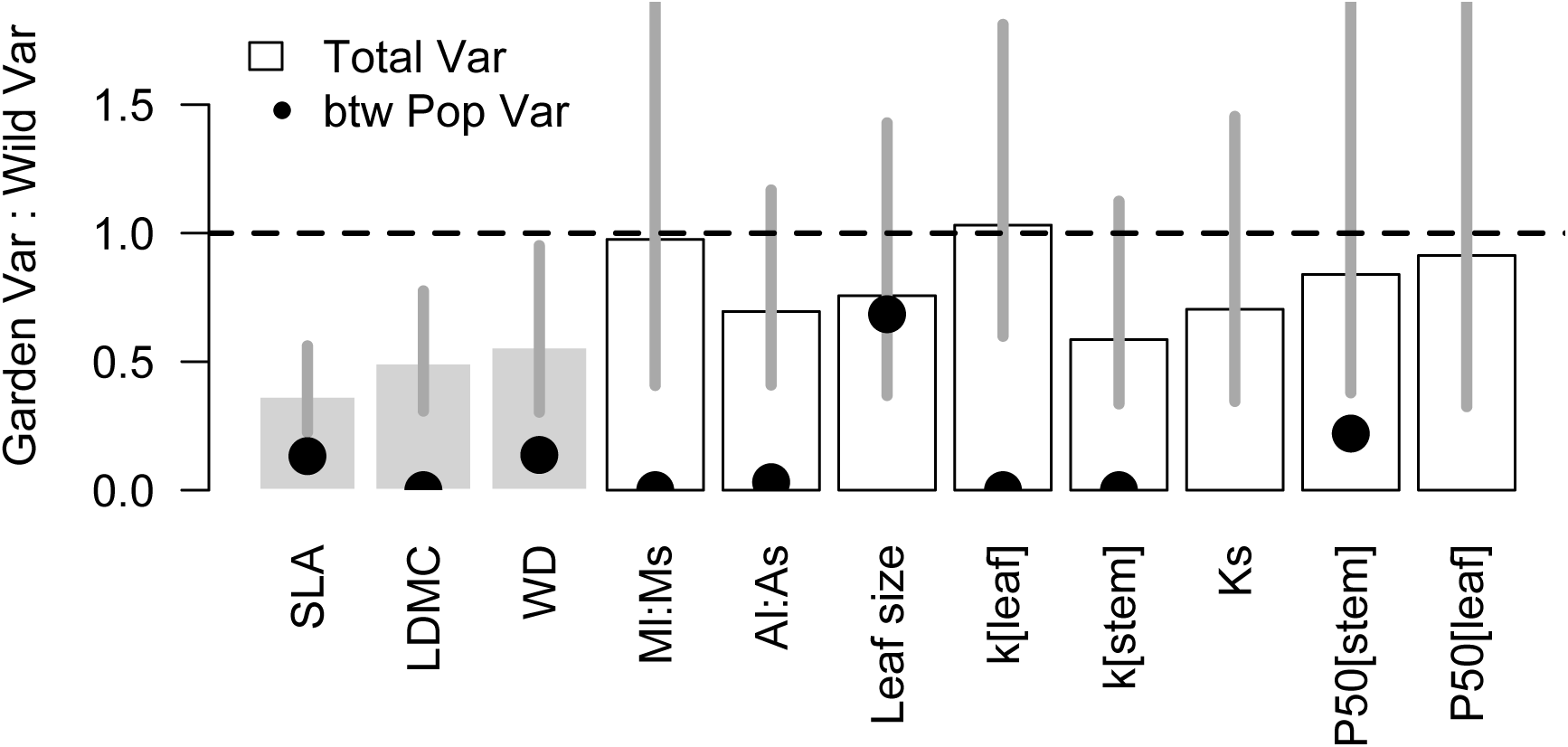
Ratio of trait variance in the common garden compared to the wild (values <1 indicating more trait variance in the wild). Bars show the ratio of total trait variance (median of 1000 bootstrapped variance estimates, lines show 5^th^ – 95^th^ percentile of ratio), gray bars indicate significantly lower variance in the garden. Points show the ratio of the between-population variance component alone (Ks and P50_leaf_ had 0 estimated between-population variance in the wild and thus no points).

Norm of reaction plots suggest that trait values and variances may be an artifact of ontogeny (younger trees in the garden than the wild) for two traits, with WD and k_leaf_ having consistently higher values in the garden than the wild for all populations (Fig. S5). For the remaining traits, genotype x environment effects appeared prevalent (i.e. reaction norms often cross). Leaf size was the only trait that did not show reaction norms of markedly opposing signs across populations. As a result, population-average trait values were entirely uncorrelated in the garden and in the wild except for M_L_:M_S_ (Fig. S2), even for the traits with similar amounts of total variation in the two settings.

### Trait coordination

Trait-trait coordination was much stronger in the wild than in the common garden, suggesting phenotypic correlations are driven by coordinated plasticity. Only six of 55 possible trait-trait correlations were significant in both the garden and the wild (Fig. 5), almost all of which are unsurprising (e.g., A_L_:A_S_ ∼ M_L_:M_S_, SLA ∼ LDMC) or mathematically related (K_s_ ∼ k_stem_ or K_s_∼A_L_:A_S_ because K_s_ was calculated as k_stem_ * A_L_:A_S_ * branch length). The main notable correlations consistent in both the garden and the wild were positive relationships between average leaf size and leaf vs stem allocation on both an area basis (A_L_:A_S_) and a mass basis (M_L_:M_S_). Moreover, average leaf size was much more strongly associated with variation in A_L_:A_S_ than number of leaves per branch in both the garden and the wild (Fig. S7), suggesting that plastic leaf expansion may be a key regulator of leaf-to-stem allocation.

**Figure 5:**
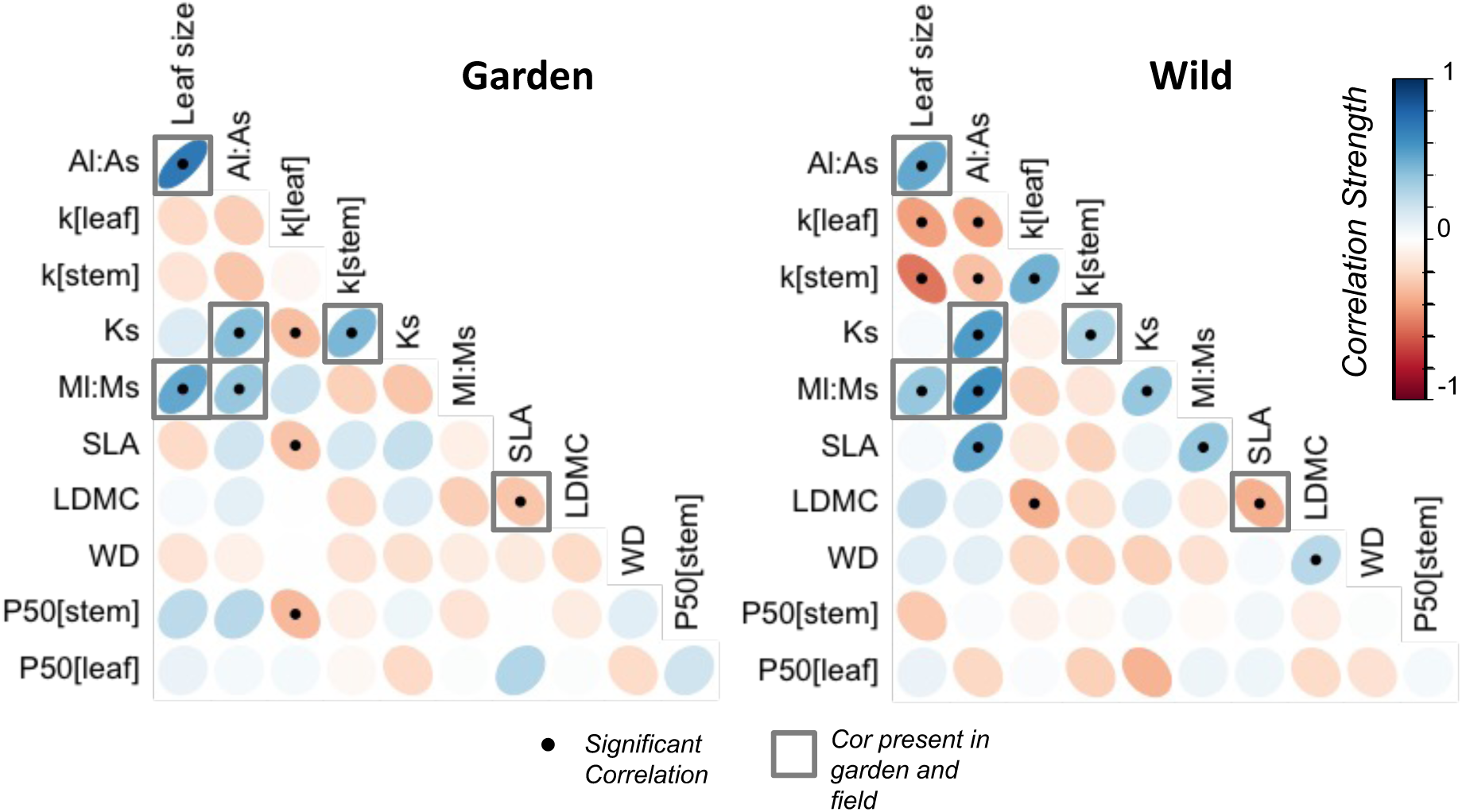
Trait integration was stronger in wild trees. Tree level trait-trait correlations in the garden (a) and in the wild (b). Ellipses and color show direction and strength of the correlation, points show statistically significant relationships, and grey boxes show relationships that are significant in both the garden and the wild.

Beyond these correlations, only three additional trait correlations were significant in the garden, while 10 additional trait correlations were significant in the wild (Fig 5). The change in trait-trait correlations from garden to wild was significant or marginally significant (α <0.05 or <0.1 respectively, based on 5000 bootstrapped correlation comparisons) in five trait pairs (Fig S8). In four of these pairs, this trend was driven by a strengthening of correlation in the wild: a increasingly negative leaf size ∼ k_stem_ correlation, and increasingly positive k_leaf_∼k_stem_; K_s_∼M_L_:M_S_ and M_L_:M_S_∼SLA correlation. Notably, hydraulic integration, namely correlations among leaf and stem conductance, leaf size and A_L_:A_S_ was strong in the wild but almost completely absent in the common garden. This plastic increase in hydraulic integration in wild trees appears to be driven by leaf size, which is strongly negatively correlated with k_leaf_ and k_stem_ in the wild (change in correlation is statistically significant for k_stem_). Across the landscape, trees with smaller leaves had lower A_L_:A_S_, but consequently greater k_leaf_ and k_stem_. We found no evidence for a tradeoff between safety and efficiency in either leaves or stems (Fig S11).

### Trait-Growth relationships

In the garden, traits were generally not strongly related to population average growth (27 yr height). LDMC was positively and k_stem_ negatively correlated with growth rates. This conflicts with expectations, if traits are presumed to drive growth (and that trait values predict the past trait values that drove growth). Leaf size and M_L_:M_S_ were the only traits correlated with growth in the expected direction, with populations with larger leaves and higher allocation to leaf mass having higher average growth rates (Fig. S9), though M_L_:M_S_ was only marginally significant.

In the wild, we found very little evidence for relationships between traits and growth (calculated as the percent of maximum size-specific Basal Area Increment), despite much larger among-population trait differentiation. The only significant relationship was a decrease in growth with increasing K_s_, which may be consistent with increasing drought avoidance in the driest populations (Fig S10). However, growth was only measured in six of the seven populations due to lost access at one site after initial trait sampling (due to threats from nearby illicit marijuana growers). Moreover, both traits and environmental conditions and resources differ among populations, making it difficult to disentangle cause versus effect.

## Discussion

We found very little among-population genetic differentiation but consistent among-population variation across the landscape in morphological and hydraulic traits of a widespread oak species. This landscape-scale variation, likely due to environmentally driven plasticity, could not be easily explained using gridded climate or annual meteorological data, but drove trait-trait coordination – particularly of leaf and stem hydraulic traits – that was otherwise lacking in the common garden.

### Traits in the common garden: little population-level differentiation

The limited evidence for genetically based trait variation that we found in a relatively mesic common garden (978 mm of mean annual precipitation, greater than all but two of the sampled populations, Table S1) contrasts with the fairly widespread experimental evidence of ecotypic variation in many trees (Leimu & Fischer 2008; Alberto *et al*. 2013), other oak species (Sáenz-Romero *et al*. 2016; Ramírez-Valiente & Cavender-Bares 2017; Browne, Wright, Fitz-Gibbon, Gugger & Sork 2019; Ramírez-Valiente *et al*. 2022), and blue oak itself (McLaughlin, pers. comm.). This contrast could potentially indicate that population-level differentiation is less common in hydraulic traits than in growth and morphological traits or that differentiation is less common in reproductively mature adult phenotypes compared to the more commonly studied seedlings/saplings. Due to the difficulty of measuring physiological traits, our sample sizes were also smaller than true quantitative genetic studies of morphological traits, and thus we way have lacked the statistical power to detect subtle among-population genetic variation. However, we note that our sample sizes are similar to or larger than prior studies that have detected among-population genetic variation in hydraulic traits (e.g. in *Corymbia callophyla*, (Blackman *et al*. 2017)). We also had sufficient power to detect significant among-population differentiation in almost all traits in wild trees. Thus, even if we missed subtle among-population genetic variation in the garden, this genetic variation must be substantially smaller than plasticity.

Among individual trees in the common garden, there was also remarkably little trait coordination, even though many of these traits are correlated across species (Mencuccini *et al*. 2019; Sanchez Martinez, Martinez-Vilalta, Dexter, Segovia & Mencuccini 2020).This lack of coordination was not necessarily an artifact of limited total trait variation in the common garden. Only three of 11 possible traits showed significantly smaller total trait variation in the garden than in the wild (Fig. 4), despite all traits showing less among-population variation in the garden. Thus, the trait variation manifested in the garden was not merely the wild trait variance minus among-site plasticity. Neither the lack of trait coordination nor the lack of correlation between wild and garden trait values (Fig. S6) was thus driven by restricted garden trait variance (i.e. limited statistical power). Moreover, there was limited evidence of populations converging on wet-adapted trait values in the garden (Fig. S5), and only WD and k_leaf_ showed offsets between the wild and garden traits that might suggest that size- or age-related effects drove garden trait values relative to wild trees. Rather, the garden trees growing in a relatively wet environment manifested a large range of phenotypes, but this variation was uncoordinated across traits and not structured by population.

At the same time, the only traits that showed expected relationships with growth rates in the common garden were leaf to stem allocation traits (leaf size and potentially M_L_:M_S_), with leafier populations showing faster growth. We conclude that changes to allometry were the primary drivers of functional differences among populations in the common garden in line with classic predictions about the drivers of growth rate variation (Lambers & Poorter 1992), and that this variation in allocation was linked in part to genetically determined variation in leaf size (Fig. S7). Meanwhile, leaf dry matter content (LDMC) and leaf-specific stem conductance (k_s_) were significantly correlated with growth rates, but in a direction contrary to expectations if lower tissue construction costs or higher hydraulic efficiency support faster growth (Hajek, Leuschner, Hertel, Delzon & Schuldt 2014). For LDMC, we interpret this as a consequence rather than a cause of growth rate variation (faster growing trees had accumulated more photosynthate in their leaves per unit leaf water by the sampling period and thus had higher LDMC), and for k_s_, results were driven by the two the two most arid populations that grew slowly in the garden but had high k_s_.

### Traits in the wild

Meanwhile, based on the wild phenotypes it appears that spatial variation in drought-related physiology is substantial in blue oak, and is likely driven by trait plasticity. In contrast to the common garden results, all traits except K_s_ and P50_stem, leaf_ showed substantial among-population trait differentiation across *Q. douglasii’*s geographic range. While all traits exhibited large branch-to-branch variation within a canopy (in both the garden and the wild), among-population trait differences were as large or larger than differences among trees in the wild (Fig 2b, 3a). Experimental evidence for plastic drought responses is reasonably common in the literature for other oak species (Gimeno *et al*. 2009; Ramirez-Valiente, Sanchez-Gomez, Aranda & Valladares 2010; Ramírez-Valiente & Cavender-Bares 2017; Ramírez Valiente, Lopez, Hipp & Aranda 2019; Ramírez-Valiente *et al*. 2022), our results highlight this plasticity as a key driver of trait variation across the landscape.

Surprisingly, this site-to-site trait variability was not strongly related to historical site climate data nor meteorology of the sampling year (absolute or anomaly relative to historical climate, Table S2, Fig. S3,S4). Even for the four traits that were significantly predicted by site climate, water limitation was not the driver of variation. Winter minimum temperatures (Tmin_gy_) was the best predictor for two traits (M_L_:M_S_ and k_stem_), perhaps indicating an early season developmental temperature constraint, and a water balance metrics (some actual evapotranspiration metric, AET_anom_ and AET_30yr_) was only a significant predictor for two traits (WD and k_leaf_, respectively). We hypothesize that these weak trait∼climate patterns are the result of *Q. douglasii*’s deep rooting strategy, which combined with the substantial geologic and edaphic variation among the sampled sites to largely decouple the actual water availability (the ‘weather underground’, sensu (McLaughlin *et al*. 2020)) from above-ground climate.

Strikingly, the plasticity-driven trait variation across California resulted in strong trait-trait coordination, particularly between hydraulic efficiency and allocation traits. Leaf and branch morphology (SLA, LDMC, WD) and P50 were largely uncoupled from other traits in both the wild and the common garden (Fig 5). However, k_leaf_ and k_stem_ significantly changed their correlation strength from uncorrelated in the garden to positively correlated in the wild, and both traits showed a strong correlation with both leaf size and A_L_:A_S_ in the wild. Collectively, this resulted in strong integration across a leaf and stem ‘hydraulic module’, wherein trees with small leaves had small A_L_:A_S_ ratios and high leaf area-specific leaf and stem hydraulic conductance.

### Leaf size-driven hydraulic coordination

In theory, the coordination or integration of plant traits across tissues should optimize whole-plant function (Reich 2014). However, across species this coordination is not universally observed among tissues (Vleminckx *et al*. 2021) or types of traits (Sanchez Martinez *et al*. 2020), and the actual mechanisms of this coordination (e.g., coordinated evolution, developmental constraints, etc.) remain elusive. The leaf and stem hydraulic integration that we found across wild populations suggests that, within individual species, plastically determined leaf size, presumably coordinated by leaf expansion, may be a key mechanism coordinating hydraulic efficiency across tissues.

In the garden we saw quite weak phenotypic trait coordination. While we are not able to directly assess genetic trait correlations (as opposed to phenotypic correlations) with this historic garden, as it was not designed for quantitative genetic analysis, our results suggest that the signal, either for genetic correlations among traits or local adaptation in these phenotypes (with the exception of leaf size) is not strong. Leaf size was the only trait that showed similar among-population variation in the wild and garden (Fig 3) and evidence of local adaptation: populations from wetter source climates grew larger leaves in the garden (Fig S3) and populations with larger average leaf sizes grew faster in the garden (Fig. S9). This hints at leaf area’s potential adaptive importance.

However, wild and garden leaf size were uncorrelated (Fig. S5,S6), suggesting strong environment and likely environment-by-genotype effects on leaf size. At the same time, wild trees showed strong hydraulic integration between the leaf area-specific hydraulic conductance of leaves and stems across wild populations. This coordination seems to be mediated by allocation to leaf area, relative to stem area (A_L_:A_S_), which was largely driven by the average size of leaves (rather than leaf number, Figure S11). Average leaf size had a larger coefficient of variation than stem diameter (0.41 versus 0.31, respectively), and had a stronger negative correlation with both k_leaf_ (ρ = −0.43) and k_stem_ (ρ= −0.54) than A_L_:A_S_ itself did with either trait (ρ= −0.39 and ρ= −0.30 for k_leaf_ and k_stem_ respectively). Further supporting the importance of plasticity, average leaf size was uncorrelated with either k_leaf_ or k_stem_ in the garden, and the increase in the correlation from garden to wild was strong for both traits and statistically significant for k_stem_ (p < 0.05). Meanwhile, the more commonly measured sapwood-specific branch conductivity (K_s_), which is standardized for branch xylem conductive area and path length, was highly variable but not differentiated among populations (Fig. 2) and unrelated to either leaf size or k_leaf_ in the wild. This may partly be due to the difficulty of measuring path length for our K_s_ estimates. In order to avoid open vessels in this long-vesseled oak species we measured the conductance of whole terminal branches cut at the petioles (rather than branch segments) and estimated the path length imperfectly from the total branch length. Thus, k_stem_ (conductance of the whole branch per unit leaf area) was likely a more accurate measurement of hydraulic efficiency than K_s_ (conductivity per sapwood area for a 1cm stem length) in this study.

Ultimately, it appears that whole-branch conductance of terminal branches and leaf conductance were coordinated in the wild not by changes in xylem efficiency itself but rather by changes in leaf size, regulated partly by ecotypic variation but largely by environmental cues governing spring leaf expansion. K_s_, or sapwood area-specific hydraulic efficiency, did not differ among populations either in the garden or the wild (Fig 1) and therefore could not drive this pattern. A developmentally fixed hydraulic architecture (in both leaves and stems) that then gets allocated to supply a variable amount of leaf area based on leaf expansion presents a parsimonious mechanism for regulating whole-tissue hydraulic resistance, as regulation of transpirational area may be much easier to regulate in response to environmental cues through turgor driven leaf expansion than anatomical changes to xylem efficiency. Moreover, the negative relationship between leaf size and k_leaf_/k_stem_ documented here contrasts with the lack of relationship between k_leaf_ and leaf size among species (Sack & Frole 2006)and suggests a developmental constraint rather than adaptive mechanism.

### Caveat to management implications

On their face, our results from mature trees might indicate that genetically informed management or restoration of blue oaks may not require strong consideration of the aridity of the seed source. Mature tree drought resilience appears to be determined by canalized physiological damage thresholds of xylem vulnerability to drought (Skelton *et al*. 2019) and largely plastic adjustments to drought avoidance-related traits (e.g. k_branch_ and k_leaf_). While we could not predict trait variation across the landscape based on site climate (Fig 2b), the substantial among-population variation in leaf-to-stem allocation and hydraulic efficiency are presumably adaptive and therefore constitute plastic adjustments to drought exposure by increasing plant hydraulic efficiency. This decreases the minimum leaf water potential required to attain a given rate of transpiration (Whitehead & Jarvis 1981; Trugman *et al*. 2019; Mencuccini *et al*. 2019) and is quite consistent with within-species adjustments to promote drought avoidance in numerous other systems (Mencuccini & Grace 1995; Martinez-Vilalta *et al*. 2009; Anderegg & HilleRisLambers 2016; Rosas *et al*. 2019; Anderegg *et al*. 2021).

Our findings of limited among-population differentiation are also consistent with the relatively small amount of local adaptation found in blue oak growth rates (Fig 1) and phenology (Papper & Ackerly 2021), and the relatively high rates of gene flow in blue oak (Papper 2021). While other California oak species have been found to show strong genetic structure across their range (Sork *et al*. 2013; Gugger, Fitz-Gibbon, Albarrán-Lara, Wright & Sork 2021), evidence suggests that blue oak population differentiation across its broad geographic distribution is more limited. Given the strong potential for plastic adjustments to plant hydraulics documented here, this might suggest that most populations can (and may) manifest more drought avoidant phenotypes in a warmer and drier future climate, regardless of their climate history.

However, we encourage caution in this interpretation, as there are two important caveats to our finding of limited drought-related among-population differentiation. First, we only studied reproductively mature trees, and our results strongly contrast with emerging results from the seedling stage that show more ecotypic differentiation (B. McLaughlin et al., 2022; B. McLaughlin in prep). Local adaptation or population-level differentiation at the earliest life stages may be the most immediately relevant for restoration, and limited differentiation in adult phenotypes does not guarantee limited differentiation in juvenile phenotypes. Indeed, explicit contrasts of among-population differentiation in adult vs juvenile phenotypes is important before we generalize across common garden results for different life stages.

Second, we were only able to perform trait measurements in a relatively wet common garden, and may not have detected important genetic differentiation in plasticity itself. If the expression of genetic differences depends on the environment – e.g., if there are strong genotype-by-environment or GxE interactions for drought-related phenotypes– then our results could differ entirely in a drier common garden location. It remains possible that genotypes from drier populations have larger potential for plasticity (as documented in a *Eucalyptus* species, Mclean *et al*. 2014), which could imply that the same source populations planted in a dry location could show substantial among-population variation due to the dry-adapted populations manifesting more plasticity than wet-adapted populations. Thus, our results do not obviate the need for proactive stewardship of genetic resources from range edge populations (McLaughlin *et al*. 2022).

## Conclusion

In a study of range-wide trait variation in blue oak (*Quercus douglasii*), we found little evidence for population-level differentiation in any of numerous morphological or hydraulic traits but substantial realized trait variation across the landscape. Our results suggest that this landscape-scale trait variation involved plastic trait adjustments, and that this plasticity drove among-tissue trait coordination. Thus, plastic adjustments, likely during the process of leaf development and expansion, drive the coordinated manifestation of drought avoidant phenotypes with high hydraulic efficiency in leaves and stems and increased carbon investment in stems relative to leaves. However, these phenotypes could not be predicted by site climate and were poorly linked to growth rates, indicating that our understanding of blue oak drought exposure and integrated performance remains limited.

## Acknowledgements

We acknowledge the Traditional Custodians and Owners of California, and recognize their continuing connection to land upon which this research was conducted. We particularly recognize the Chumash, Sierra Miwok, Konkow, Pomo, Northern Wintu, on whose traditional territory our labs and study sites sit. We thank the Hopland Research Extension Center for access to the common garden and keeping the trees alive for multiple decades and the Ranger Station at Sonora for access to trees on their property. We thank CALFIRE for saving the Hopland common garden trees from the Mendocino Complex Fire. We also thank S. Mazer for insightful comments on the analysis. We acknowledge funding from the National Science Foundations (NSF 1457400 to DDA & TED; NSF DBI-1711243, 2003205, and 2216855 to LDLA) and the National Oceanographic and Atmospheric Administration (Climate and Global Change Fellowship to LDLA).

## Data Availability

The data from this study are publicly available in Dryad at [insert Dryad DOI] (data citation). All code to replicate the statistical analyses and generate the paper figures are in the Github repository https://github.com/leanderegg/BlueOakGarden.git.

## Author contributions

LDLA, RPS, PP, DDA and TED conceived of the project ideas; LDLA, RPS and PP designed the methodology; LDLA, RPS and JD collected the data; LDLA analyzed the data and lead the writing of the manuscript. All authors contributed to editing the manuscript.

## Notes

https://github.com/leanderegg/BlueOakGarden.git

### Competing Interest Statement

The authors have declared no competing interest.

### Summary of Updates

Introduction and Discussion updated to clarify importance and limitations (i.e. inferences about plasticity) of the study, additional analyses to included in the Supplemental files

