## Supplementary material for "Plasticity, geographic variation and trait coordination in blue oak drought physiology": Table S1

**Supplemental Information for: Plasticity, geographic variation and trait coordination in blue oak drought physiology**

**Table S1: Site characteristics for the seven populations used in the study.** 30-year climate normals (1951-1980) were extracted from the Basin Characterization Model (Flint *et al.* 2013). Tmin = mean temperature of the coldest month, Tmax = mean temperature of the warmest month, PPT= total precipitation, PET = potential evapotranspiration, CWD = climatic water deficit (potential evapotranspiration – modeled actual evapotranspiration).

| Site Name | Lat (°N) | Lon (° W) | Elev (m) | Tmin (°C) | Tmax (°C) | PPT (mm) | PET (mm) | CWD (mm) |
|---|---|---|---|---|---|---|---|---|
| **LYN** | 40.34686 | 121.8794 | 658 | 7.3 | 22.1 | 849.1 | 1144.4 | 671.7 |
| **SJR** | 37.09536 | 119.7387 | 342 | 7.8 | 24.5 | 471.7 | 1367.3 | 993.3 |
| **SMR** | 35.08696 | 120.0659 | 610 | 8.6 | 23.1 | 560.5 | 1351.9 | 985.8 |
| **SON** | 37.81632 | 120.6546 | 104 | 7.9 | 23.5 | 454.7 | 1284.7 | 981.7 |
| **SRD** | 37.98214 | 120.3677 | 638 | 6.2 | 22.2 | 836.6 | 1264.5 | 783.3 |
| **SUG** | 38.07000 | 120.2200 | 1020 | 4.9 | 21.2 | 1014.3 | 1193.3 | 733.9 |
| **SMT** | 40.66595 | 122.3721 | 230 | 10.4 | 23.6 | 1221.9 | 1174.5 | 690.1 |
| **Hopland Garden** | 39.01721 | 123.0909 | 275 | 6.1 | 22.8 | 977.7 | 1169.1 | 732.6 |

**Table S2: Statistical results of trait ANOVAs and linear relationships with climate in the common garden and the wild.** Traits were averaged to the individual.

| Location | Trait | n | eta2 | omega2 | ANOVA p-value | Pop Rand Effect for trait~clim | Best Climate Predictor | Delta AIC | Clim p-value | Clim R2 |
|---|---|---|---|---|---|---|---|---|---|---|
| Garden | SLA | 35 | **0.36** | **0.22** | **0.039** | no | **PET[30yr]** | **-9.6** | **8.00E-04** | **0.29** |
| | LDMC | 35 | 0.18 | 0 | 0.43 | no | none | 0 | 1 | 0 |
| | WD | 34 | 0.24 | 0.07 | 0.237 | no | **PPT[30yr]** | **-1.86** | **0.0467** | **0.12** |
| | ml_ms | 35 | 0.12 | -0.06 | 0.693 | no | none | 0 | 1 | 0 |
| | Al_As | 37 | 0.27 | 0.13 | 0.115 | no | none | 0 | 1 | 0 |
| | leafsize | 35 | **0.4** | **0.27** | **0.017** | no | **PPT[30yr]** | **-8.11** | **0.0018** | **0.26** |
| | kleaf | 35 | 0.14 | -0.04 | 0.591 | no | Tmin[30yr] | -0.28 | 0.1147 | 0.07 |
| | kstem | 35 | 0.21 | 0.04 | 0.325 | no | PET[30yr] | -0.05 | 0.1319 | 0.07 |
| | Ks | 35 | 0.13 | -0.05 | 0.636 | no | none | 0 | 1 | 0 |
| | P50stem | 33 | 0.21 | 0.02 | 0.379 | no | none | 0 | 1 | 0 |
| | P50leaf | 18 | 0.54 | 0.27 | 0.134 | no | none | 0 | 1 | 0 |
| | Growth | 453 | **0.2** | **0.17** | **0** | yes | **PET[30yr]** | **-5.75** | **0.0081** | **0.05** |
| Wild | SLA | 47 | **0.6** | **0.54** | **0** | **yes** | AET[gy] | -0.3 | 0.3531 | 0.08 |
| | LDMC | 46 | **0.55** | **0.48** | **0** | **yes** | none | 0 | 1 | 0 |
| | WD | 51 | **0.42** | **0.33** | **0** | **yes** | **AET[anom]** | **-5.35** | **0.0249** | **0.27** |
| | ml_ms | 45 | **0.48** | **0.4** | **0** | **no** | Tmin[gy] | -15.14 | 0 | 0.32 |
| | Al_As | 48 | **0.55** | **0.48** | **0** | **yes** | CWD[anom] | -1.18 | 0.136 | 0.21 |
| | leafsize | 48 | **0.39** | **0.3** | **0.002** | **yes** | Tmin[gy] | -1.04 | 0.1538 | 0.14 |
| | kleaf | 33 | **0.52** | **0.4** | **0.003** | **yes** | CWD[gy] | -1.82 | 0.0877 | 0.22 |
| | kstem | 33 | **0.51** | **0.39** | **0.003** | **yes** | **Tmin[gy]** | **-3.43** | **0.0482** | **0.28** |
| | Ks | 33 | 0.06 | -0.15 | 0.932 | no | none | 0 | 1 | 0 |
| | P50stem | 37 | 0.2 | 0.04 | 0.301 | no | **AET[30yr]** | **-3.74** | **0.017** | **0.15** |
| | P50leaf | 23 | 0.09 | -0.24 | 0.943 | no | none | 0 | 1 | 0 |
| | Growth | 32 | **0.44** | **0.33** | **0.007** | **yes** | none | 0 | 1 | 0 |

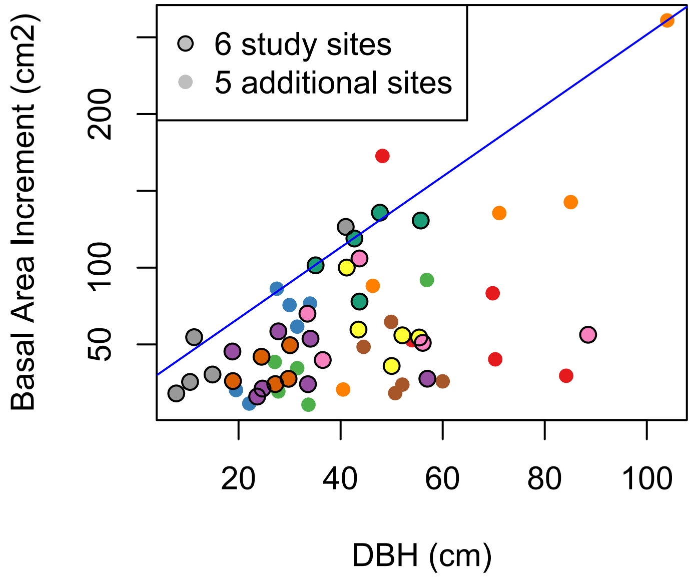


**Figure S1:** *Total 5-year Basal Area Increment (2013-2018) measured from tree cores versus tree DBH. Raw 5yr BAI was standardized for tree size by calculating the '% of max BAI', or the % of observed BAI compared to the 90$^{th}$ quantile regression of BAI versus DBH for the tree's specific DBH (blue line). Points outlined in black indicate trees from the 6 sites in this study (one site was dropped due to lost access). Points without black outlines indicate trees from an additional 5 sites from across California, USA. Colors indicate sites.*

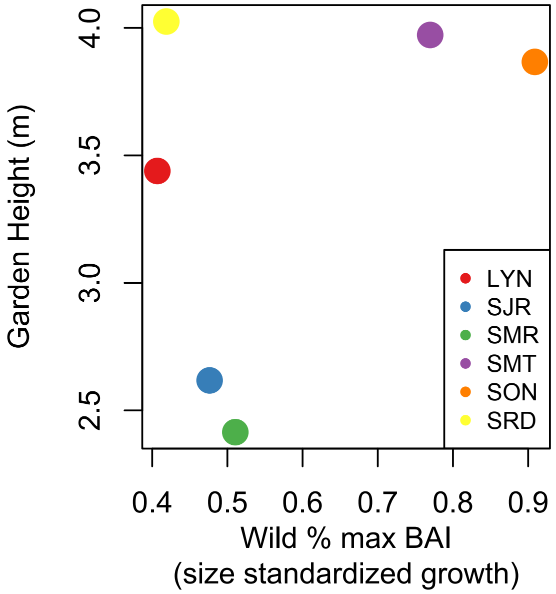


**Figure S2:** *Growth in the garden was uncorrelated with growth in the wild source populations, showing little evidence of transgenerational carryover effects (e.g. stressed parent trees resulting in low garden performance).*

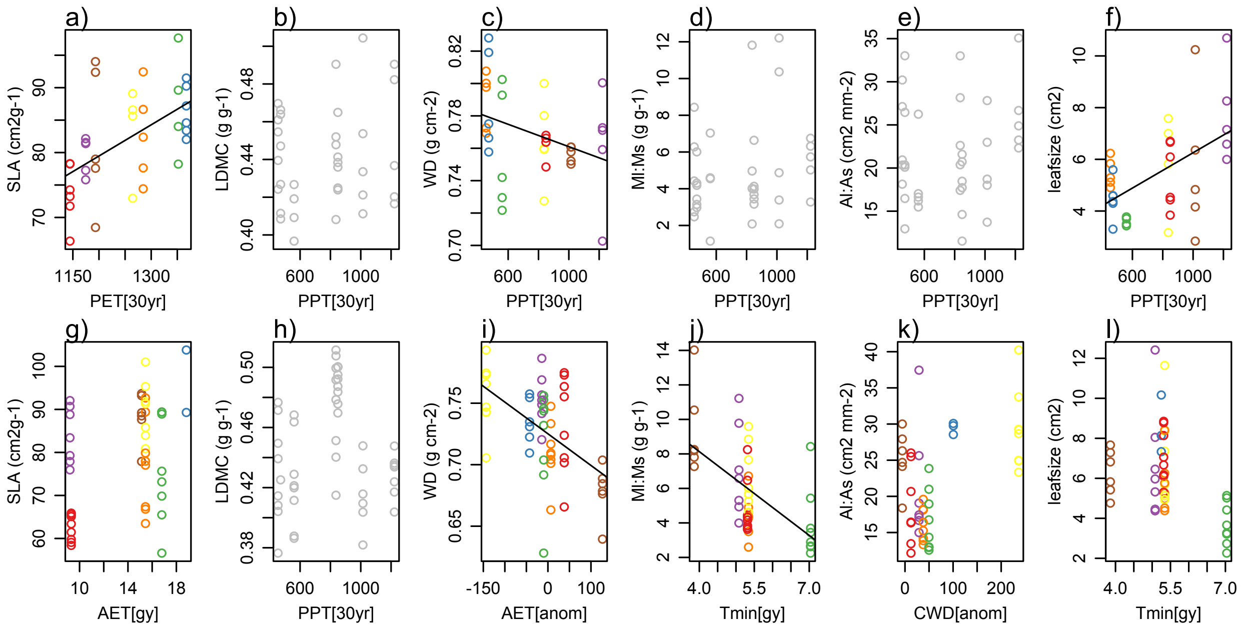


**Figure S3:** *Relationships between morphological or allocation traits (averaged to the individual) and source population climate (garden measurements, top panels) or site climate (wild measurements, bottom panels). Traits are plotted against the climate variable found to be the best predictor based on AICc, except for traits plotted in gray for which no climate variable had a lower AICc than the null model. These traits are plotted against 30year mean annual precipitation (PPT[30yr]). Solid trend lines indicate significant relationships ($p<0.05$), dashed trend lines indicate marginal significance ($p<0.1$), and no trend lines indicate that the climate variable had a lower AICc than the null model but was not significant. Colors indicate site, as in Fig S2 and Fig 1. PET: potential evapotranspiration, PPT: precipitation, AET: actual evapotranspiration, CWD: climatic water deficit, Tmin: minimum annual temperature. [30yr] indicates 30 year climate mean, [gy] indicates the weather conditions from the 2017-18 water year/growth year, [anom] indicates the difference between gy conditions and 30yr mean conditions.*

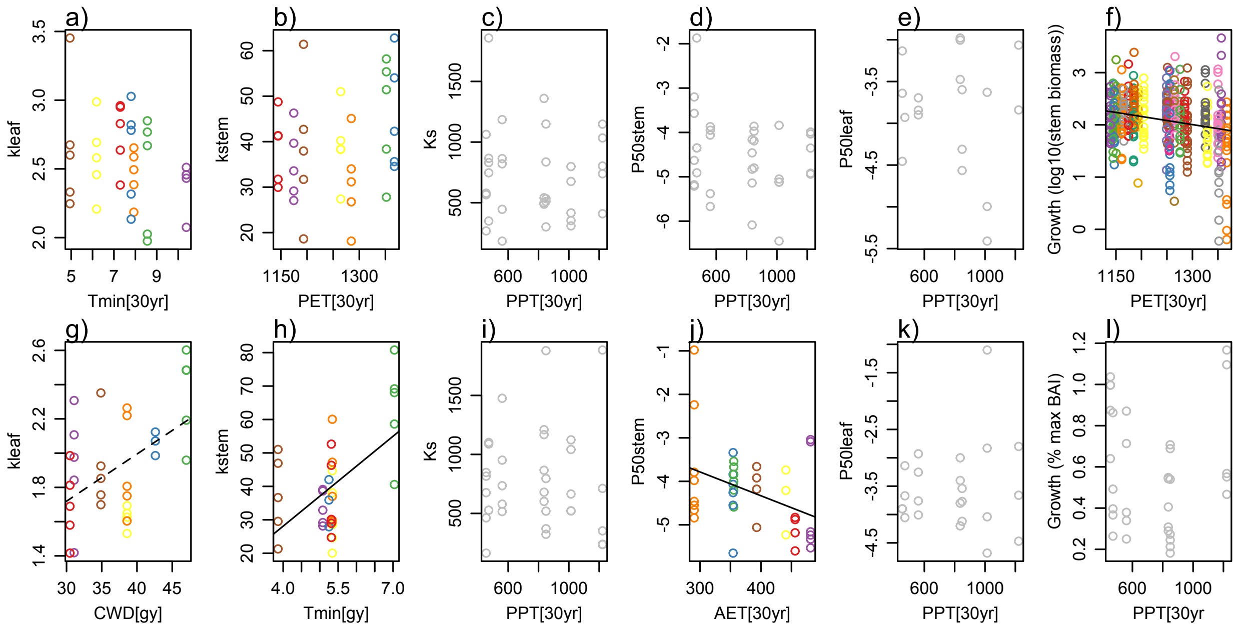


**Figure S4:** *Relationships between hydraulic traits and tree growth (averaged to the individual) and source population climate (garden measurements, top panels) or site climate (wild measurements, bottom panels). Traits are plotted against the climate variable found to be the best predictor based on AICc, except for traits plotted in gray for which no climate variable had a lower AICc than the null model. These traits are plotted against 30year mean annual precipitation (PPT[30yr]). Solid trend lines indicate significant relationships (p<0.05), dashed trend lines indicate marginal significance (p<0.1), and no trend lines indicate that the climate variable had a lower AICc than the null model but was not significant. PET: potential evapotranspiration, PPT: precipitation, AET: actual evapotranspiration, CWD: climatic water deficit, Tmin: minimum annual temperature*

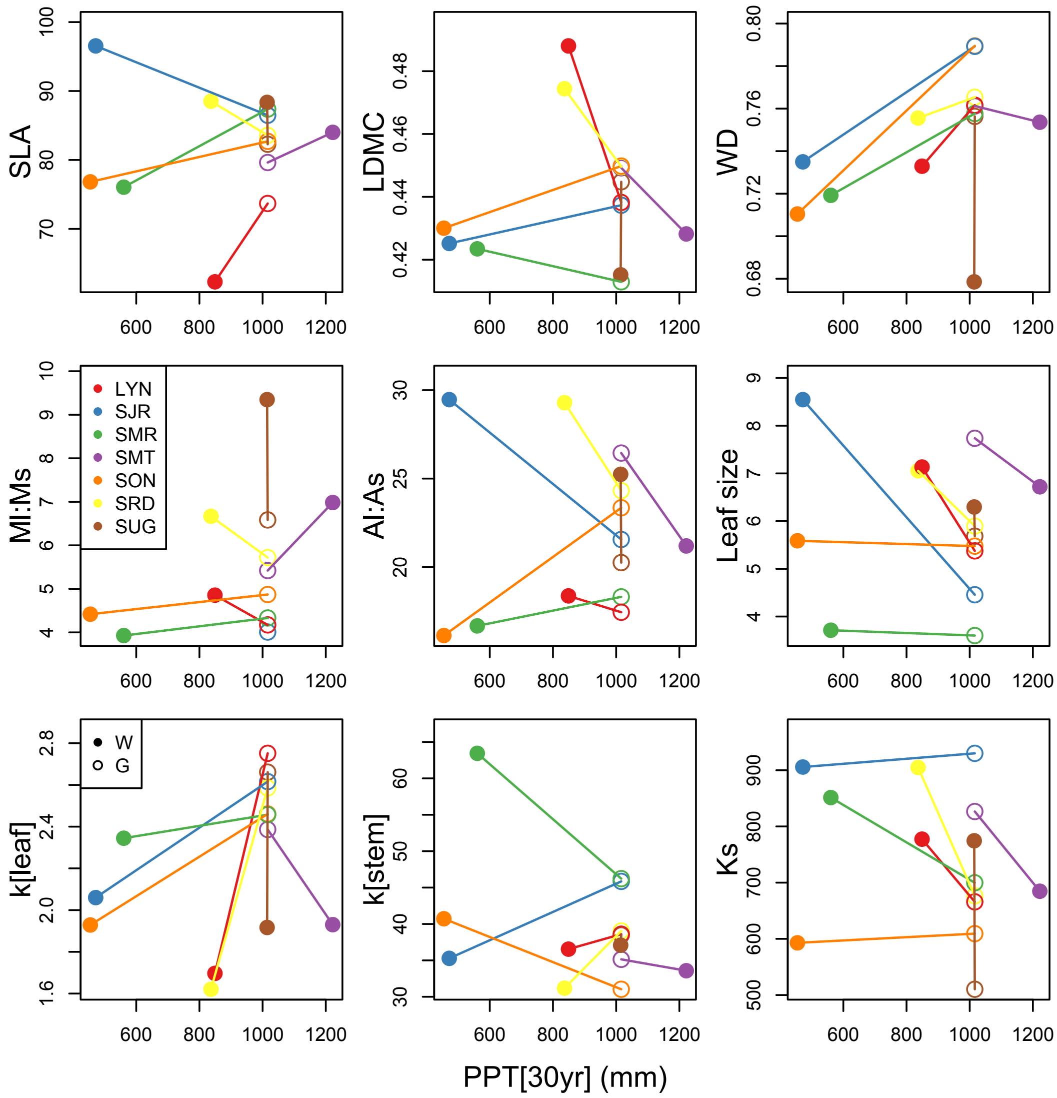


***Figure S5:*** *Norm of reaction plots showing population mean trait change from the Wild to the Garden, based on 30yr mean annual precipitation (PPT[30yr]). Results are qualitatively similar regardless of climate metric (e.g. sampling year meteorology, 30 year normal of various water balance metrics).*

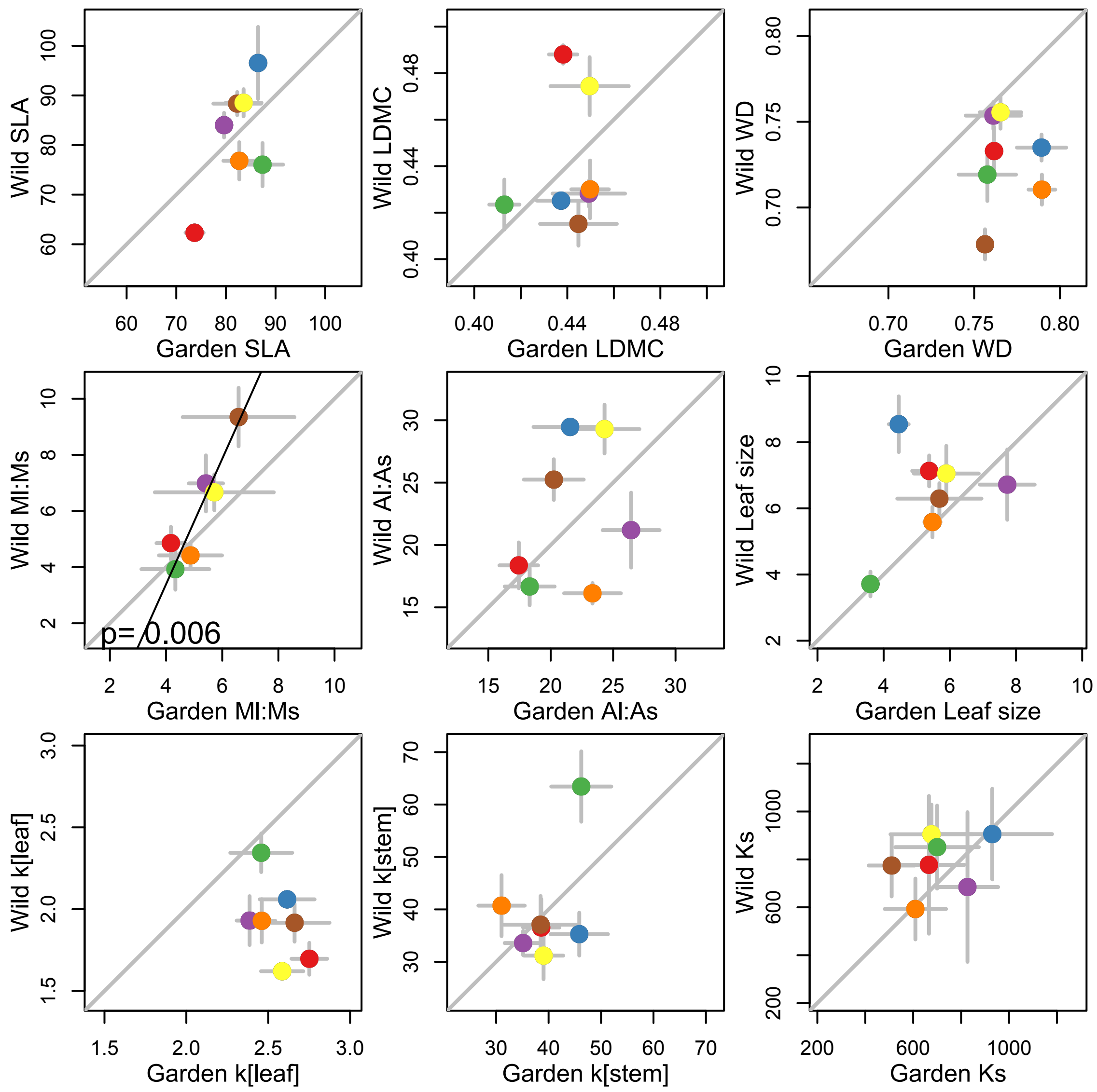


**Figure S6:** *Population mean trait values in the garden are not correlated with population mean trait values in the wild for any trait. Points show population means, error bars show population standard errors. Gray line shows 1:1 relationship. Population colors as in Fig. 1 and Fig. S2*

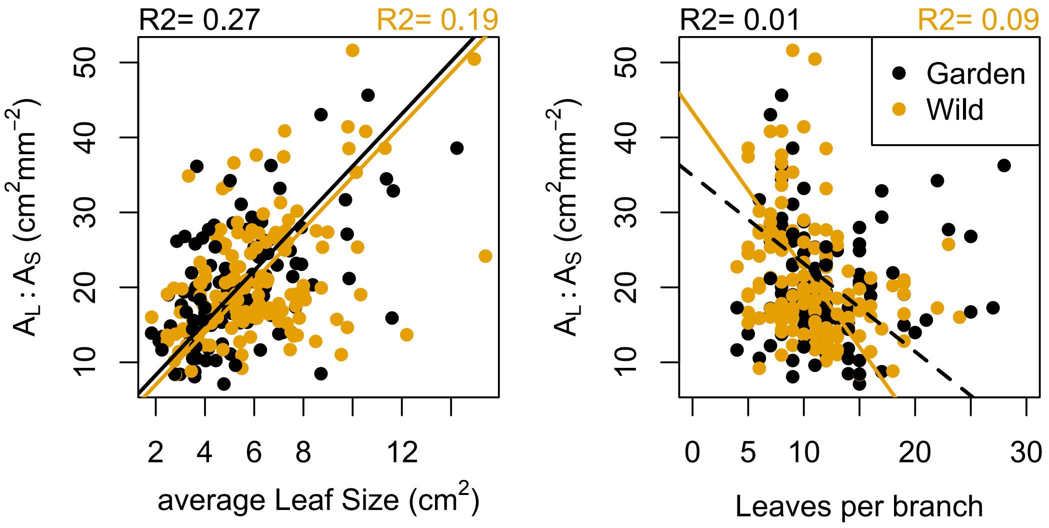


**Figure S7:** *Average leaf size per branch is a stronger driver of terminal branch $A_L$:$A_S$ than number of leaves per branch.*

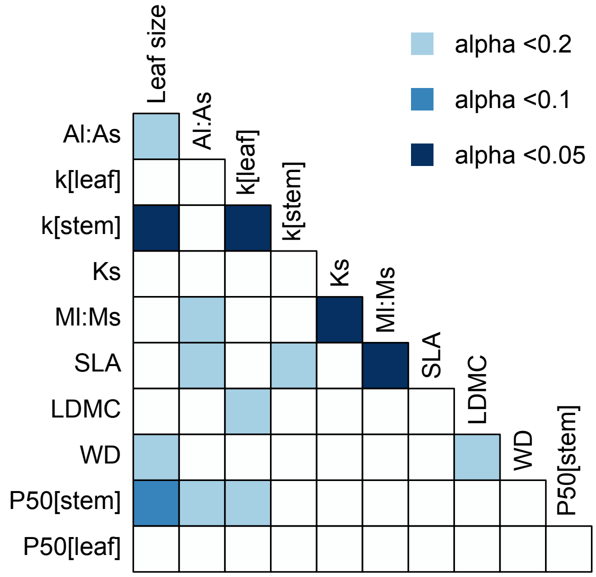


**Figure S8:** *Multiple trait correlations were significantly different in the wild versus the garden (based on 5000 bootstrapped comparisons of correlations), particularly k[stem] relationships relating to leaf size and k[leaf].*

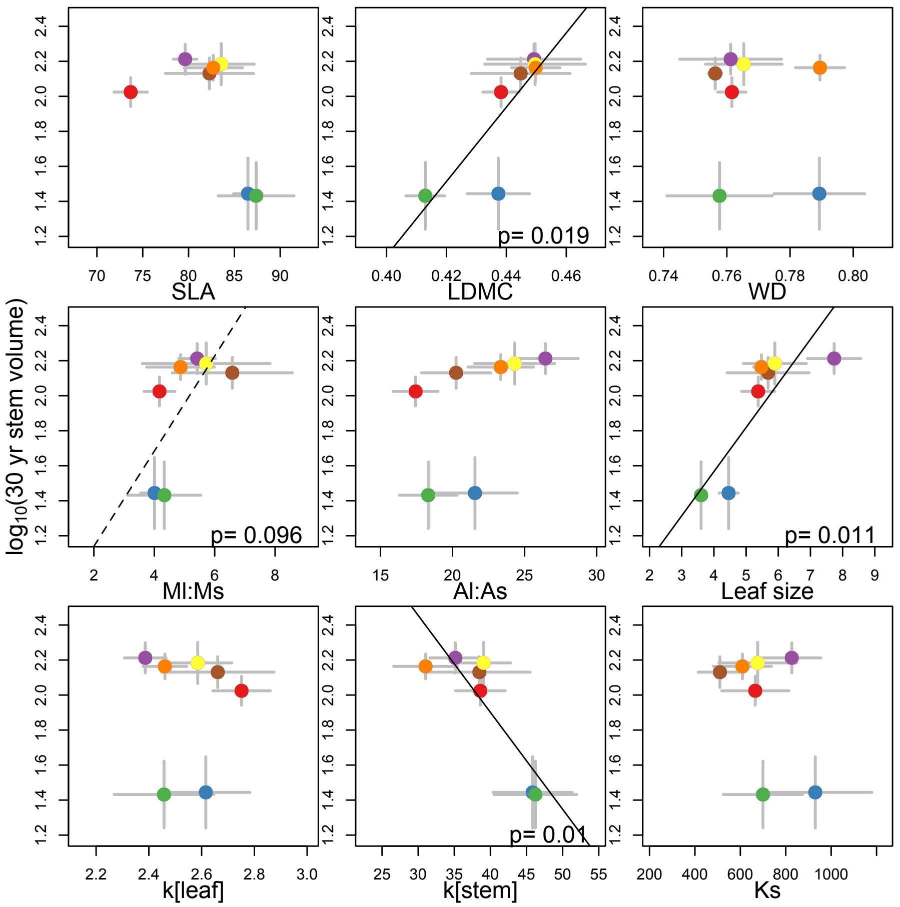


**Figure S9:** *In a mesic common garden, hydraulic traits rarely relate to average growth rates as expected. Panels show population average trait values and population mean height attained after 30 years of growth (+/- 1 se error bars). Site colors as in Fig. 1/S2.*

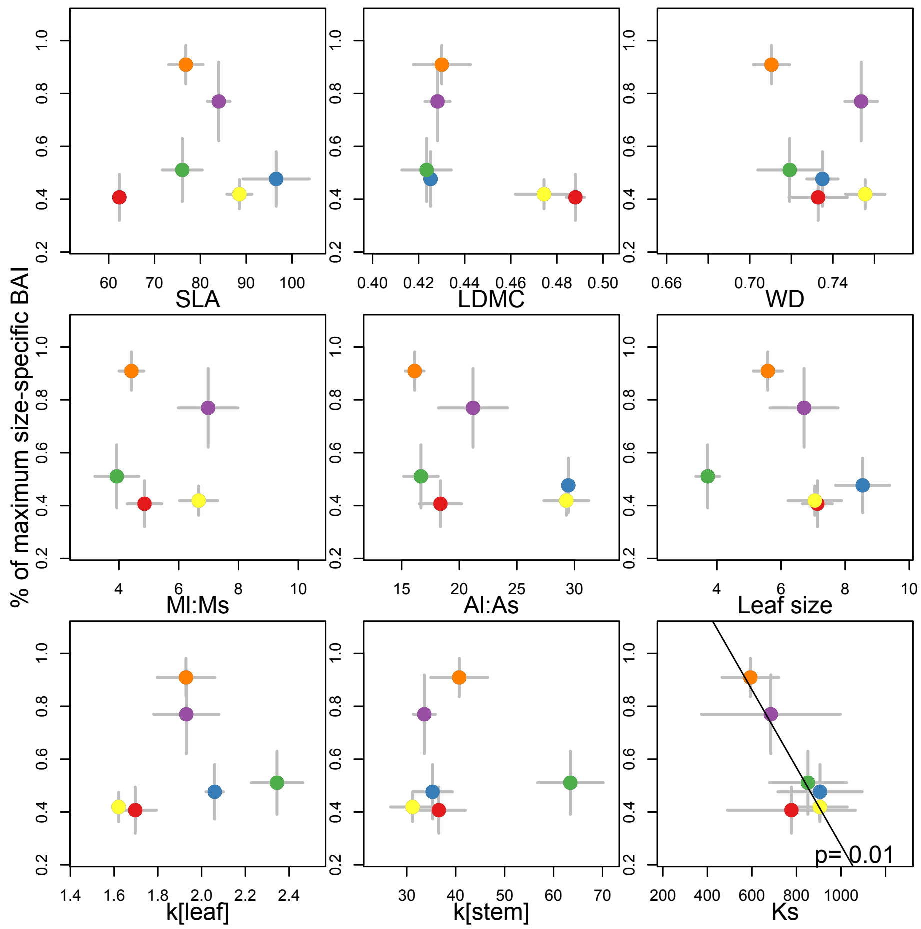


**Figure S10:** *In the wild, with the exception of $K_s$, no population average traits are correlated with population average growth rates. $K_s$ shows a negative correlation, with populations with higher sapwood-specific hydraulic efficiency having slower growth rates. Growth rates were calculated as the average % of maximum Basal Area Increment (BAI) given a tree's DBH. Growth is only available for six of the seven sampled populations because we could not revisit the seventh to collect cores (got run off by pot growers). Site colors as in Fig. 1/S2.*

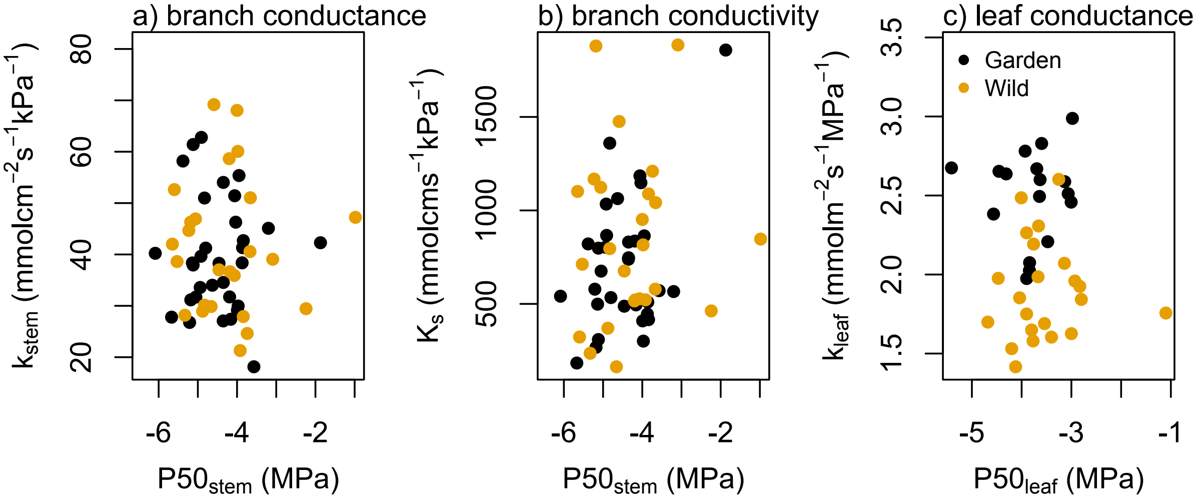


**Figure S11:** *We found no evidence for a 'safety versus efficiency' tradeoff among trees, either in the garden or in the wild. Stem leaf-specific conductance (a) and sapwood area-specific conductivity (b) show no significant relationship with stem P50, nor does leaf conductance with leaf P50 (c).*
